# *C. elegans* DBL-1/BMP Regulates Lipid Accumulation via Interaction with Insulin Signaling

**DOI:** 10.1101/191049

**Authors:** JF Clark, M Meade, G Ranepura, DH Hall, C Savage-Dunn

## Abstract

Metabolic homeostasis is coordinately controlled by diverse inputs, which must be understood to combat metabolic disorders. Here we introduce DBL-1, the *C. elegans* BMP2/4 homolog, as a significant regulator of lipid homeostasis. We used neutral lipid staining and a lipid droplet marker to demonstrate that both increases and decreases in DBL-1/BMP signaling result in reduced lipid stores and lipid droplet count. We find that lipid droplet size, however, correlates positively with the level of DBL 1/BMP signaling. Regulation of lipid accumulation in the intestine occurs through non-cell-autonomous signaling, since expression of SMA-3, a Smad signal transducer, in the epidermis (hypodermis) is sufficient to rescue the loss of lipid accumulation. Finally, genetic evidence indicates that DBL-1/BMP functions upstream of Insulin/IGF-1 Signaling (IIS) in lipid metabolism. We conclude that BMP signaling regulates lipid metabolism in *C. elegans* through inter-organ signaling to IIS, shedding light on a less well-studied regulatory mechanism for metabolic homeostasis.

## Introduction

Metabolic homeostasis in animals relies on integrating input from numerous sources to coordinately regulate energy intake and expenditure across multiple organs. Nutrient sensing pathways relay signals in the absence or presence of food. Simultaneously, feedback loops are triggered in response to the lack or abundance of cellular resources. Satiety responses are produced to slow the intake of energy when sufficient resources have been obtained. The balance of these pathways provides the basis of metabolic homeostasis. Metabolic disorders, such as type II diabetes, are the result of an imbalance in the signaling web that is homeostasis. The last few decades have provided numerous insights into this regulatory network, but new regulatory interactions continue to be discovered.

The Transforming Growth Factor beta (TGFβ) superfamily is a major group of peptide ligands conserved across animal phyla. The superfamily includes the founding TGFβs, as well as Bone Morphogenetic Proteins (BMPs), growth and differentiation factors (GDFs), Activin, Nodal, and others. These peptides signal through conserved signal transduction pathways responsible for development, growth, and differentiation (Shi and Massague 2003; Wu and Hill 2009). Intriguingly, emerging evidence from correlative studies in humans, as well as in vivo studies in mice, implicate several TGFβ ligands in lipid metabolism and homeostasis (Wang et al. 2004; Fain et al. 2005; Sjoholm et al. 2006; Bottcher et al. 2009; Shen et al. 2009).

BMPs are a group of TGFβ-related signals with key regulatory roles in development and differentiation. Mammalian BMP2 and BMP4 play important roles in early development and cell differentiation, as well as being critical for bone and cartilage development (Chen et al. 2004). In Drosophila, the BMP ligand DPP is required for dorsoventral patterning of the early embryo as well as later patterning of the imaginal discs (Spencer et al. 1982; Padgett et al. 1987). However, the homeostatic roles of BMP ligands are less well studied. We have used molecular genetic and imaging tools available in the nematode *C. elegans* to gain insight into the homeostatic roles of BMP ligands.

*C. elegans* has a conserved BMP signaling pathway that includes founding members of the Smad family of signal transducers, SMA-2, SMA-3, and SMA-4 (Savage et al. 1996). DBL-1, the *C. elegans* BMP2/4 homolog, plays a major role in body size regulation, male-tail development, and mesodermal patterning (Suzuki et al. 1999; Foehr et al. 2006). Initial evidence of a role for DBL-1 in metabolism came from our microarray analysis of genes regulated by the DBL-1 pathway. This analysis identified several genes related to fat metabolism including genes encoding fatty acid desaturases and genes involved in β-oxidation (Liang et al. 2007). *C. elegans* is a prominent model system for the study of lipid homeostasis, and is particularly suitable for the identification of cell and tissue interactions that mediate homeostasis (Ashrafi 2007). Although nematodes do not possess dedicated adipocytes, they store triglycerides in lipid droplets in the intestine and in epidermal tissue (the hypodermis) via biochemical mechanisms that are evolutionarily conserved.

In this study, we show that DBL-1/BMP signaling plays an important role in regulating lipid stores in *C. elegans*. Alterations to DBL-1 signaling levels result in a loss of lipids and changes in lipid droplet morphology. DBL-1 signaling acts non-cell-autonomously in the hypodermis to regulate lipid storage in the intestine. Finally, we show that the lipid phenotype of *dbl-1* mutants is reliant on Insulin/IGF-1-like Signaling (IIS), a well-known regulator of fat metabolism (Kimura et al. 1997).

## Results

### DBL-1/BMP Signaling is Required for Lipid Accumulation

Our previous data identified numerous lipid metabolism genes as downstream targets of DBL-1/BMP signaling via microarray analysis (Liang et al. 2007). These genes include *fat-6* and *fat-7*, which encode Δ9 fatty acid desaturases homologous to stearoyl coA desaturase (SCD). Mutations in either gene result in an overall decrease in neutral lipids, in addition to other phenotypes (Watts and Browse 2002; Ntambi et al. 2004). To determine if DBL-1 signaling is important for lipid metabolism, we measured stored lipid levels in animals with altered DBL-1 function as well as those with mutations in *fat-6, fat-7*, or both. Animals were grown at the standard temperature of 20°C and stained with Oil Red O, at the fourth larval stage (L4), to quantify overall neutral lipid content, including lipid storage in the intestine and in the hypodermis. Consistent with previous reports, we observed a significant decrease in Oil Red O staining in *fat-6, fat-7*, and *fat-6;fat-7* mutant animals by 45.2%, 46.1%, and 47.0%, respectively, compared to wild type. Similarly, we observed a decrease in *dbl-1* mutants by 34.8% compared to wild type. This decrease was not significantly different from that observed in the *fat* mutants (p>0.38). We also analyzed a loss of function mutation in *lon-2*, which encodes a negative regulator of DBL-1 signaling (Gumienny et al. 2007). Interestingly, *lon-2* mutants also had a reduction in staining, at levels similar to that of *dbl-1* mutants (p=.99) (Figure 1A, B). A possible cause for the decrease in lipid storage is a reduction in food uptake, as seen in *eat-2* mutants (Raizen et al. 1995). However, *dbl-1* mutants had a very slight increase in the rate of pharyngeal contractions, while *sma-3* animals had no significant change from wild type (Figure S1), and thus a decrease in food uptake is not likely to explain the decrease in lipid storage in *dbl-1* or *sma-3* mutants.

**Figure 1.**
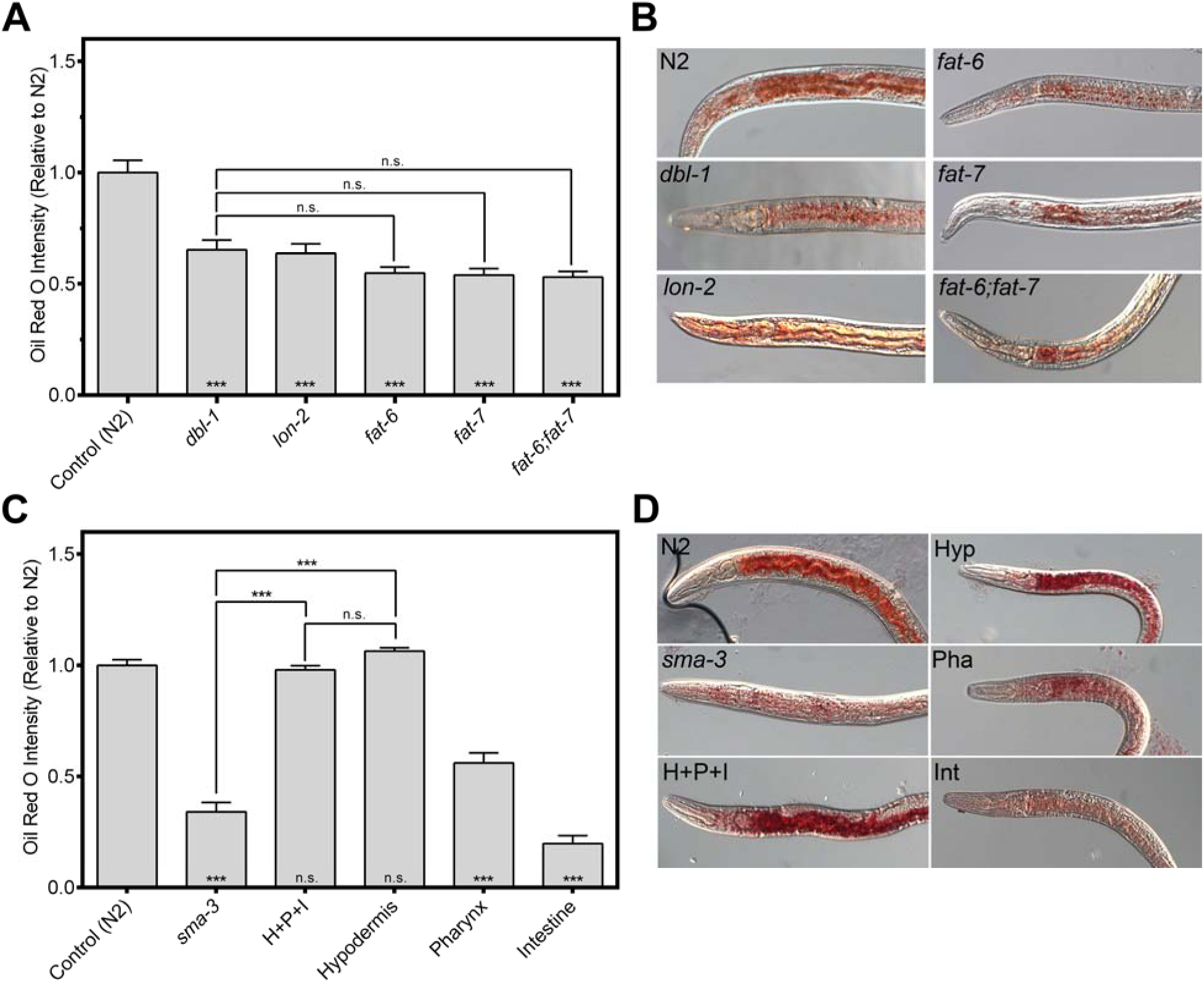
Functions and Tissue-Specificity of DBL-1 Signaling in Lipid Storage. A) Both *dbl-1* and *lon-2* mutants show a decrease in lipid levels via Oil Red O staining, similar to that of *fat* mutants. Animals were grown at 20°C and stained at the L4 larval stage. B) Images of L4 animals stained with Oil Red O taken at 400X. C) Loss of *sma-3* results in decreased lipid levels. Expression of *sma-3* in the hypodermis is sufficient to rescue the lipid phenotype. Expression in either the pharynx or the intestine cannot rescue lipids to wild-type levels. Animals were grown at 20°C and stained at the L4 larval stage. D) Images of L4 animals stained with Oil Red O taken at 400X. Anterior is to the left. For all graphs, asterisks across the bottom denote significance compared to Control, n.s. not significant, *** p value < .001, error bars denote SEM.

### Hypodermal Expression of Smads is Required for Lipid Accumulation

This lipid storage phenotype led us to question in which tissues DBL-1 signaling is necessary to regulate fat metabolism. *C. elegans* do not have dedicated adipocytes, so fat is stored in the intestine and in the hypodermis. DBL-1 signaling components (receptors and Smads) are expressed in the pharynx, hypodermis, and intestine. We have previously shown that expression of *sma-3/Smad* in the hypodermis is sufficient and necessary to rescue the small body size phenotype of *sma-3* mutants (Wang et al. 2002). We took the same approach of directing expression of *sma-3(+*) to specific tissues, in an otherwise *sma-3* mutant background and measured fat accumulation using Oil Red O.

Surprisingly, *sma-3* expression in the hypodermis was also required for rescue of the low-fat phenotype in *sma-3* mutants (Figure 1C, D). Although *sma-3* expression in the pharynx resulted in a slight increase in Oil Red O staining, intestinal *sma-3* expression in *sma-3* mutants, resulted in a further decrease in staining, compared to *sma-3* mutant animals. Thus, expression in either pharyngeal tissue or intestine was not sufficient to rescue the lipid stores in *sma-3* mutants to wild-type levels. However, when *sma-3* was expressed in either the hypodermis or all three tissues, Oil Red O intensity was rescued to wild-type levels with no significant difference observed (p>0.78) (Figure 1C, D). These data indicate that Smad activity functions non-autonomously in the hypodermis to regulate fat storage in the intestine.

### DBL-1/BMP Functions Upstream of IIS to Regulate Lipid Accumulation

Our tissue specific rescue experiments suggest that SMA-3/Smad activity in the hypodermis may regulate the expression of a secreted ligand that signals to the intestine. One potential mechanism by which this signaling may occur is through the regulation of insulin ligands. *C. elegans* uses multiple insulin-like ligands that act through a single insulin receptor, DAF-2/InsR (Pierce et al. 2001). The IIS pathway in *C. elegans* was first identified for its role in regulating dauer development. A temperature sensitive mutation in daf-2, when exposed to the restrictive temperature of 25°C, results in the development of *daf-2/InsR* mutants into dauers, an alternate L3 stage utilized for survival in high stress environments. In addition, *daf-2* mutants exhibit other phenotypes, including increased lifespan and stress tolerance (Gottlieb and Ruvkun 1994; Tissenbaum and Ruvkun 1998). Moreover, the involvement of Insulin/IGF-1-like Signaling (IIS) in fat metabolism is well documented in *C. elegans* (Kimura et al. 1997). Notably, our microarray analysis revealed *ins-4* as a transcriptional target of the DBL-1 pathway; expression was increased in both *dbl-1* and *sma-9* mutant backgrounds. INS-4 is an insulin-like ligand expressed in the hypodermis, in addition to neurons (Ritter et al. 2013).

We therefore constructed strains containing mutations in both BMP and IIS pathways to test interactions between these pathways via genetic epistasis. To bypass dauer arrest in *daf-2* mutant animals, these experiments were conducted by raising animals at the permissive temperature of 15°C followed by a shift to 25°C. Our findings confirmed the high fat phenotype of *daf-2* mutants, exhibiting an average increase of 17.7% over wild type. Similarly, *daf-2;sma-3* double mutants showed an average increase in Oil Red O intensity of 11.4% (Figure 2A), which is not significantly different from *daf-2* (p=0.71). We concluded that the *daf-2* high-fat phenotype is epistatic to the *dbl-1* low-fat phenotype, suggesting that *daf-2* acts downstream of *dbl-1* with regard to fat storage regulation.

**Figure 2.**
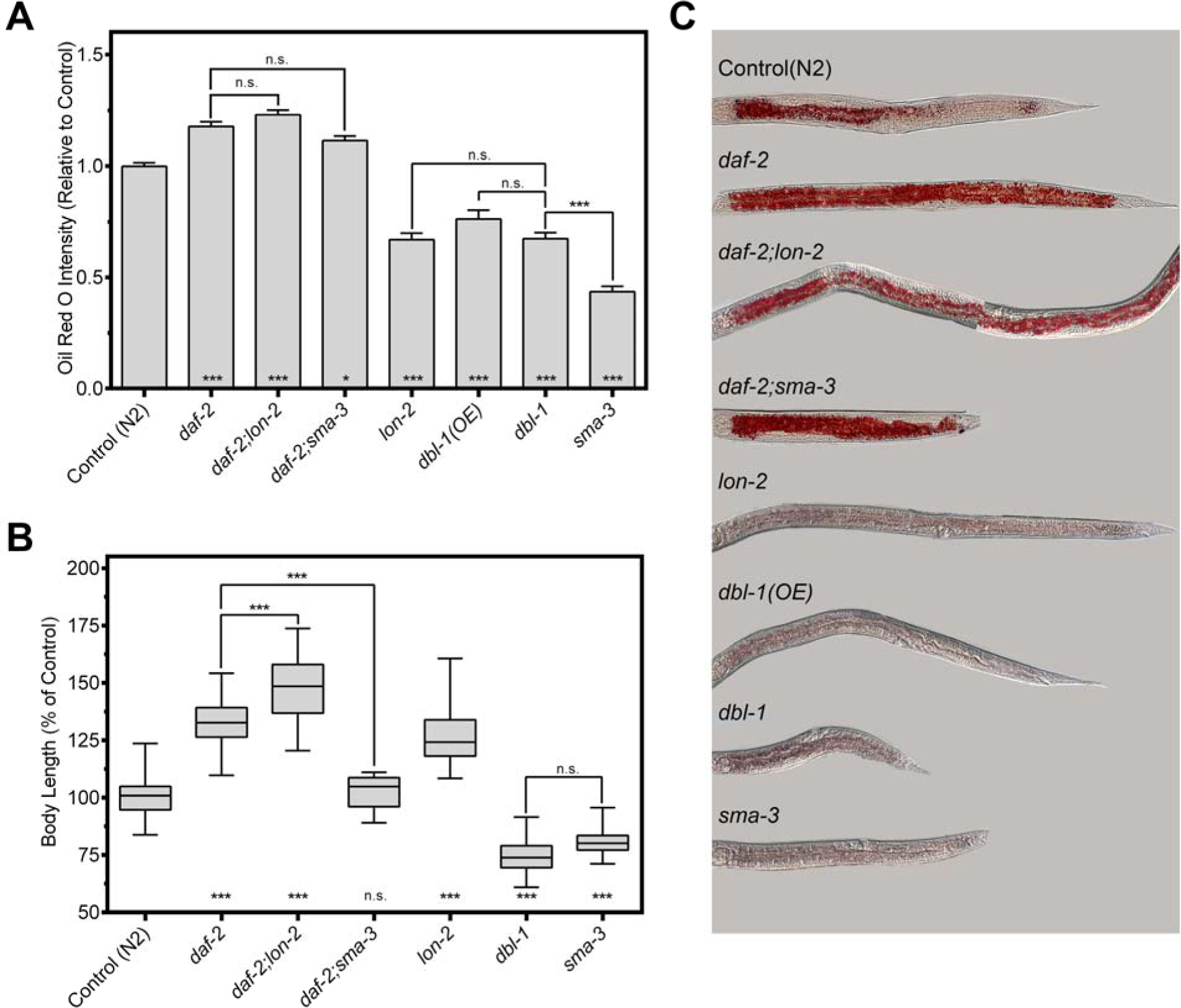
DBL-1 Signaling Functions Upstream of IIS to Regulate Lipid Storage but not Body Size. A) *daf-2;lon-2* and *daf-2;dbl-1* display the high fat phenotype of *daf-2* animals using Oil Red O. Epistasis analysis places DBL-1 signaling upstream of daf-2/InsR. *dbl-1(OE)* animals show a similar decrease in lipid levels as *dbl-1* and *lon-2* mutants. Animals were grown at 15°C until the L2/L3 molt and then shifted to 25°C and stained at the L4 larval stage. Error bars denote SEM. B) Animals containing mutations in both DBL-1 pathway components and *daf-2* exhibit additive body sizes. Both pathways regulate body size independently. Animals were grown at 15°C until the L2/L3 molt and then shifted to 25°C and measured at the L4 larval stage. Boxes denote the 2^nd^ and 3^rd^ quartiles, whiskers denote min and max values. C) Images of L4 animals stained with Oil Red O taken at 200X. Anterior is to the left. For all graphs, asterisks across the bottom denote significance compared to Control, n.s. not significant, * p value < .05, *** p value < .001.

We also assessed mutants in other components of the DBL-1 pathway. *sma-3* mutants had an average reduction in Oil Red O intensity of 56.5%. To examine increased DBL-1 activity, we used both a *dbl-1* overexpression strain and *lon-2* mutants, which are deficient in a negative regulator of DBL-1. Consistent with the observation from *lon-2* mutants, a *dbl-1* overexpression strain [*dbl-1(OE)*] also exhibited a reduction in lipid stores by an average of 23.9% (Figure 2A, C). This finding indicates precise regulation of DBL-1 signaling is required to maintain wild-type levels of lipids. It also indicates that the lipid and body size phenotypes of the DBL-1 pathway are independent, as both long and short animals exhibited decreased lipids. *daf-2;lon-2* double mutants, like *daf-2;dbl-1* mutants, have an increase in fat accumulation indistinguishable from that of *daf-2* single mutants (p>0.71). This epistatic relationship between the two pathways suggests that IIS functions downstream of DBL-1 signaling in the regulation of fat accumulation.

### DBL-1/BMP and IIS Pathways Independently Contribute to Body Size

We also tested whether these two pathways interact to regulate body size. Small body size was the first identified phenotype of the DBL-1 pathway (Savage et al. 1996; Suzuki et al. 1999). We measured animals at the L4 stage, benchmarked against wild-type controls: *dbl-1* mutant animals showed an average body length reduction of 25.3%; *sma-3* body length was reduced by 19.2%. Conversely, an increase in DBL-1 signaling resulted in an increase in body size, as *lon-2* mutants exhibited an average increase of 26.2%. Interestingly, a downregulation of DAF-2/InsR showed a significant increase in body size by an average of 31.9% compared to wild type. When animals with mutations in both pathways were measured, an additive effect was observed. The *daf-2;lon-2* mutant animals had an average increase in length of 48.0% compared to wild type, while the *daf-2;sma-3* mutant animals were comparable to the length of wild type (Figure 2B, C). These data are consistent with an independent study by the Baumeister lab (Qi et al. 2017) and suggests that the DBL-1 and IIS pathways work independently to regulate the overall body size of *C. elegans*.

### Changes to DBL-1/BMP Signaling Levels Alter Lipid Droplet Morphology

To determine how the DBL-1 pathway regulates lipid stores at a subcellular level, we analyzed lipid droplet morphology. Lipid droplets are vital for the utilization and storage of energy at the cellular level and are highly regulated. The morphology of lipid droplets can be an indicator of the function and well-being of a cell, or an organism. In humans, the function of white adipose tissue and brown adipose tissue differ significantly; these differences primarily reflect how they regulate lipid droplets (Meex et al. 2009; Yu et al. 2015; Luo and Liu 2016). We first examined lipid droplet morphology via electron microscopy. Images of *dbl-1, sma-3*, and *sma-9* mutants depict worms with much smaller lipid droplets in the intestine, compared to that of wild-type animals (Figure 3).

**Figure 3.**
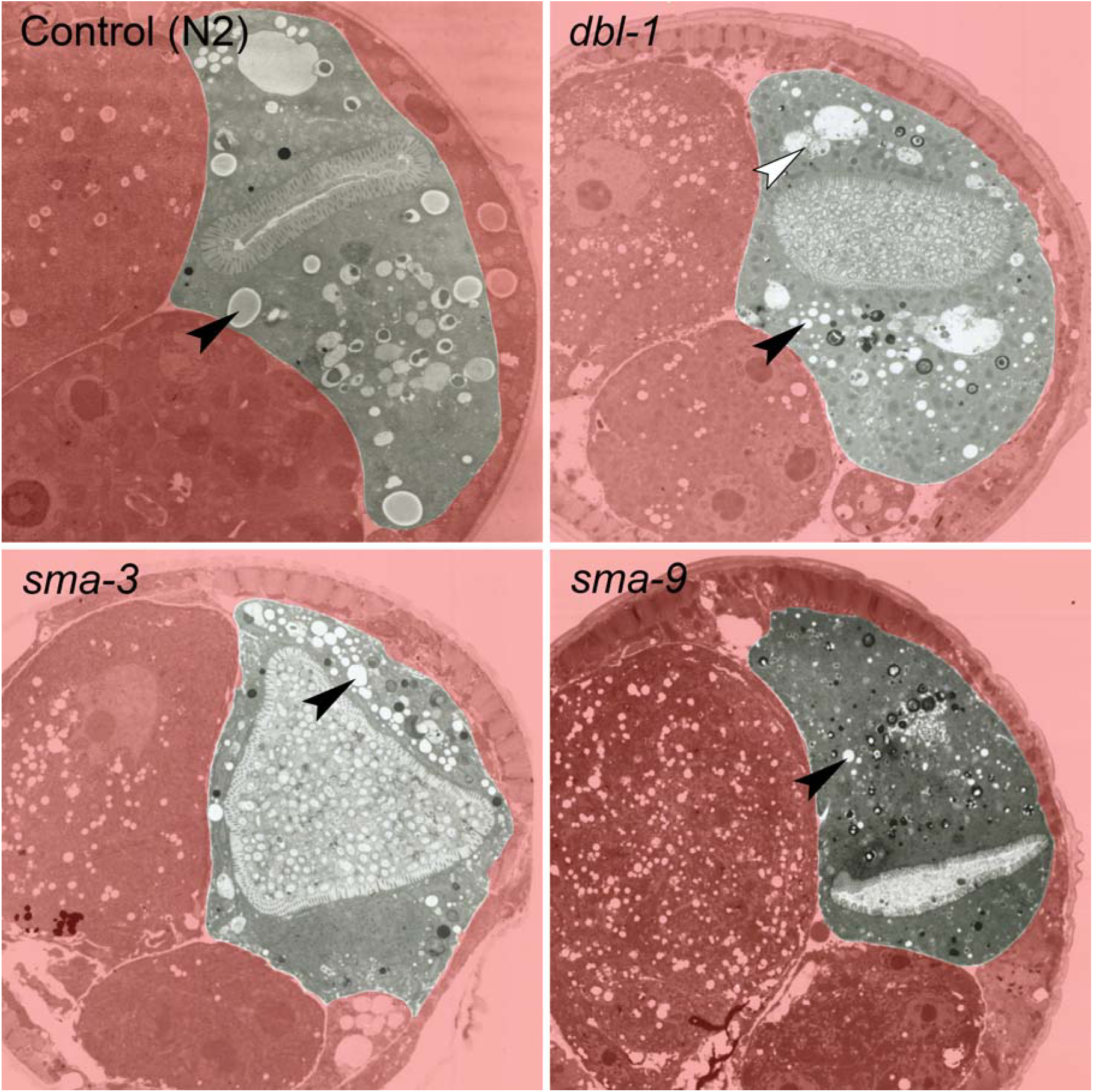
DBL-1 Pathway Mutants Exhibit a Decrease in Lipid Droplet Size via Electron Microscopy. *dbl-1, sma-3*, and *sma-9* mutants mid-body sections display smaller lipid droplets compared to those seen in wild-type. Black arrowheads depict representative lipid droplets, white arrowhead depicts lysosome-like organelle. Animals were grown at 20°C and imaged at adulthood 2 days post egg lay.

To visualize lipid droplets in a larger sample size, we used transgenic animals expressing DHS-3::GFP (Figure 4). DHS-3 is a short-chain dehydrogenase shown to bind to the surface of lipid droplets (Zhang et al. 2010). DHS-3::GFP is expressed specifically in the intestine and not in the hypodermis. Animals were grown and analyzed at the standard growth temperature of 20°C. Our data indicate that lipid droplet size is positively correlated with DBL-1 signaling levels. Wild-type animals with DHS-3::GFP have an average lipid droplet diameter of 1.02 μm, with a maximum diameter of 2.26 μm (Figure 4A, C). The *dbl-1* mutants show a decrease in average droplet diameter to 0.58 μm (p<0.001), with a maximum diameter of only 1.06 <m, a decrease of about 53%. The overexpression strain shows an increase to 1.13 μm on average (p<0.001), with a maximum diameter of 2.54 <m, an increase of over 12%. The *dbl-1* mutants displayed the lowest level of variance indicating that a decrease in *dbl-1* expression confines the size of lipid droplets to a smaller range compared to wild type. The *dbl-1(OE*) mutants displayed a slightly larger level of variance compared to wild type, which indicates an increase in the maximum size of lipid droplets, but does not restrict the minimum size compared to wild type.

**Figure 4.**
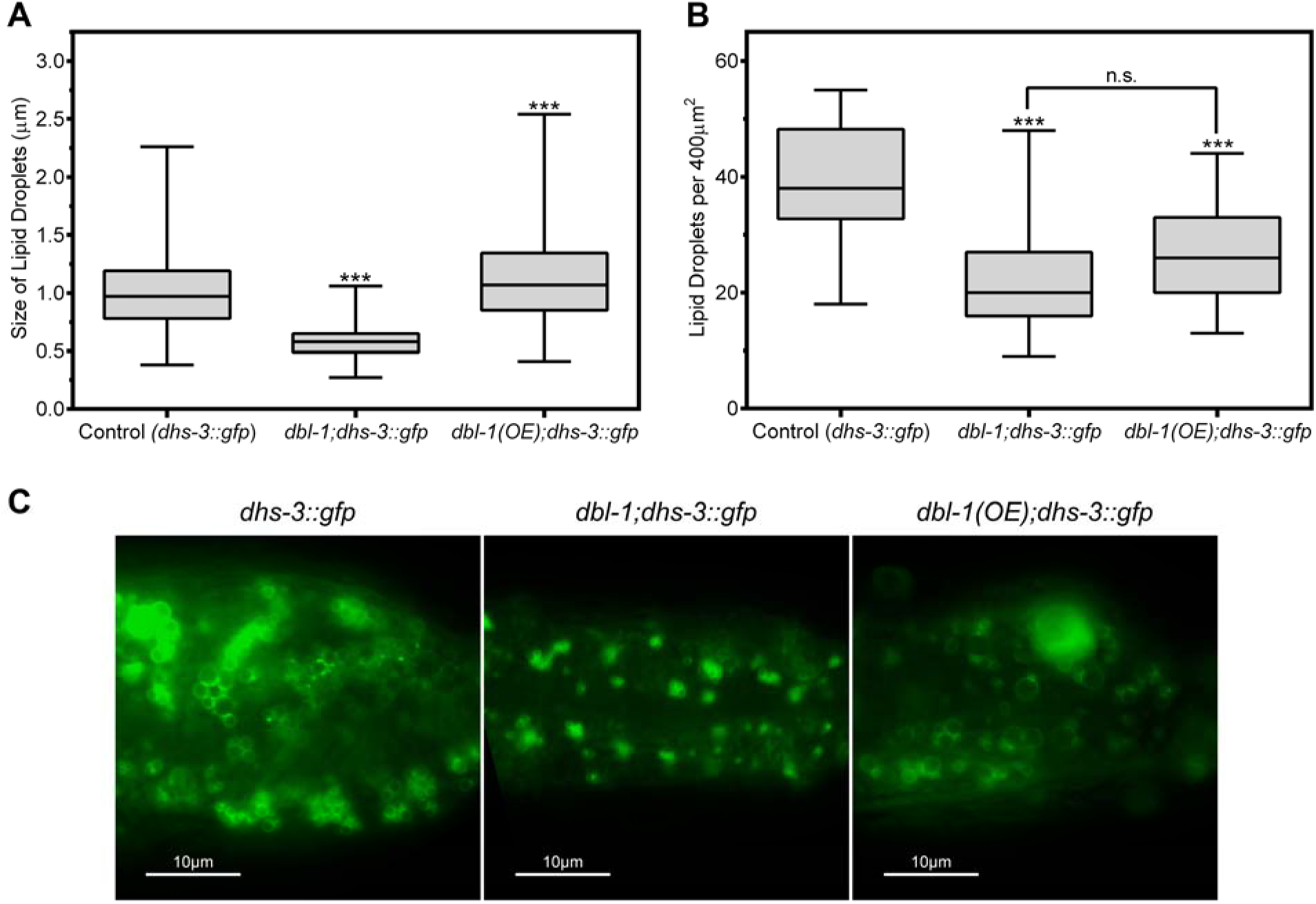
DBL-1 Pathway Regulates Lipid Droplet Morphology. A) Lack of *dbl-1* results in significantly smaller lipid droplets, while overexpression of *dbl-1* increases the average diameter of lipid droplets. *dhs-3::gfp* was used to visualize lipid droplets. Animals were grown at 20°C and imaged at the L4 larval stage. B) Both a loss or an increase in DBL-1 signaling results in a reduction in the overall number of lipid droplets in the animals. *dhs-3::gfp* was used to visualize lipid droplets. Animals were grown at 20°C and imaged at the L4 larval stage. C) Images of L4 animals with altered levels of DBL-1. Images depict the posterior end of the intestine taken at 1000X. For all graphs, n.s. not significant, *** p value < .001, boxes denote 2^nd^ and 3^rd^ quartiles, whiskers denote min and max values.

However, the trend was different when comparing the average number of lipid droplets per 400 μm^2^. Wild-type animals had an average of 39.6 droplets, while both *dbl-1* and *dbl-1(OE*) showed a decrease, 23.8 (p<0.001) and 26.7 (p<0.001), respectively (Figure 4B, C). The *dbl-1(OE*) mutants displayed a lower level of variance compared to the wild-type and *dbl-1* strains, which may help account for their overall decrease in lipids observed via Oil Red O staining (Figure 2A). Thus, while both decreasing and increasing DBL-1 signaling resulted in a reduction in total fat accumulation, the mechanisms at the level of lipid droplet size are different.

### DBL-1/BMP Regulation of Lipid Droplet Morphology is Partially Dependent on IIS

Next, we wanted to determine if IIS is required for the lipid droplet phenotypes of DBL-1 signaling mutants. To bypass dauer arrest in *daf-2* mutant animals, these experiments were conducted by raising animals at the permissive temperature of 15°C followed by a shift to 25°C. In the *daf-2* mutant background, the average diameter of lipid droplets was 1.12 μm with a maximum diameter of 2.63 μm. This was significantly larger than those of wild type with an average of 0.90 μm and a maximum diameter of 2.37 μm (p<0.001) (Figure 5A, C). The *daf-2* mutants also had the greatest level of variance, indicative of having a greater range of diameters than any of the other strains. The *daf-2;dbl-1* double mutants had an average diameter of 0.97 μm and a maximum diameter of 2.34 μm. This size was similar to that of *daf-2* mutants, but still slightly smaller than in *daf-2* mutants (p<0.001). Thus, for lipid droplet size, we do not observe the strict epistatic relationship seen in the Oil Red O experiments. These data suggest that the interaction between IIS and DBL-1 signaling may be slightly more complex and nonlinear at the lipid droplet level, with the possibility of a DAF-2-independent function for DBL-1. Alternatively, the deviation from epistasis may be due to our use of *daf-2(e1370*), a strong loss-of-function but not a null allele, which would be inviable.

Next, we examined the average number of lipid droplets in the animals. *daf-2* exhibited a significant increase compared to wild type, 54.5 and 34.0 (p<0.001), respectively. The *daf-2;dbl-1* animals were similarly increased over wild type, at 46.1 droplets (p=0.039). The increased count of *daf-2* and *daf-2;dbl-1* were similar to each other and not significantly different (p=0.276). Interestingly, the *dbl-1* mutant did not exhibit a significant decrease in lipid droplet count compared to wild type, 26.6 versus 34.0 (p=0.471); the variance between the two samples was also very similar, suggesting no change between the two strains. These data suggest that DBL-1 regulation of lipid droplet count may be temperature sensitive (Figure 5B, C).

**Figure 5.**
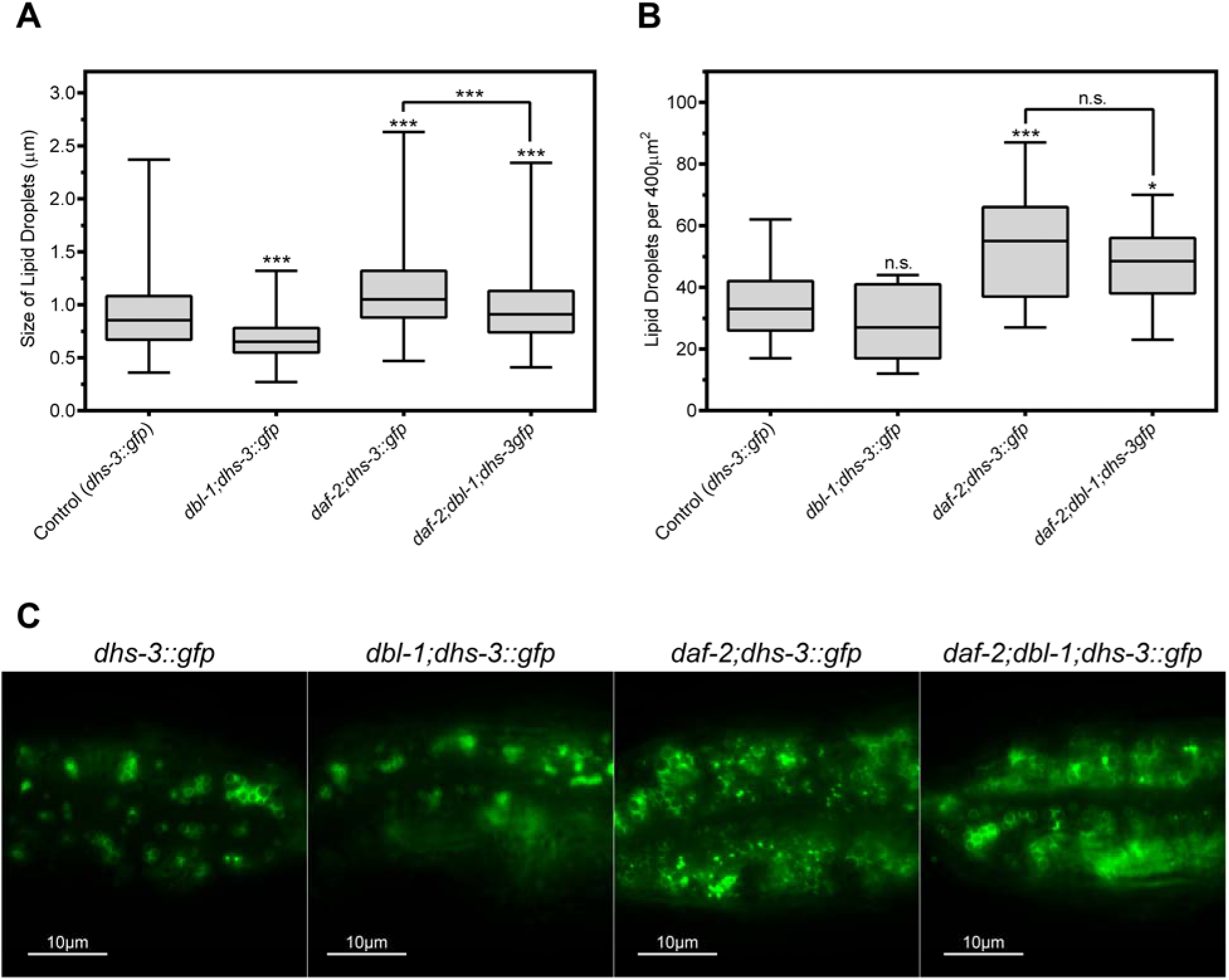
DBL-1 Regulates Lipid Droplet Size in part via IIS. A) The average diameter of lipid droplets in *daf-2;dbl-1* mutants is similar to but slightly distinct from that of *daf-2* single mutants, suggesting a partially independent effect from the DBL-1 pathway. *dhs-3::gfp* was used to visualize lipid droplets. Animals were grown at 15°C until the L2/L3 molt and then shifted to 25°C and imaged at the L4 larval stage. B) *daf-2* mutants exhibit an increase in lipid droplet count compared to both wild-type and *dbl-1*. At 25°C, *dbl-1* does not have a significant effect on the number of lipid droplets. *dhs-3::gfp* was used to visualize lipid droplets. Animals were grown at 15°C until the L2/L3 molt and then shifted to 25°C and imaged at the L4 larval stage. C) Images of L4 animals containing mutations in either *daf-2, dbl-1*, or both. Images depict the posterior end of the intestine taken at 1000X. For all graphs, n.s. not significant, * p value < .05, *** p value < .001, boxes denote 2^nd^ and 3^rd^ quartiles, whiskers denote min and max values.

## Discussion

In this study, we have shown for the first time that DBL-1/BMP signaling plays an important role in lipid metabolism, as DBL-1 pathway mutants, as well as *dbl-1* overexpression, have a distinct decrease in lipid stores via Oil Red O staining. In addition, lipid droplet morphology of DBL-1 pathway mutants is significantly changed from wild type. We have identified a non-autonomous mechanism for this regulation, since DBL-1 signaling activity is required in the hypodermis to maintain proper lipid levels in the intestine. The epistasis of *daf-2/InsR* in *daf-2;dbl-1* double mutants indicate that DBL-1 regulates lipid levels by modulation of DAF-2/InsR activity. These data implicate a BMP - IIS signaling axis as a major player in the regulation of lipid metabolism. Our model is illustrated in Figure 6. DBL-1 is expressed primarily in neurons, from which it signals to the Smad pathway in the hypodermis, leading to inhibition of the IIS pathway and stimulation of lipid storage. Observed differences in the lipid droplet morphology between *daf-2;dbl-1* double mutants and *daf-2* single mutants suggest that there may also be DAF-2-independent regulation of lipids by DBL-1 (arrow with question mark).

**Figure 6.**
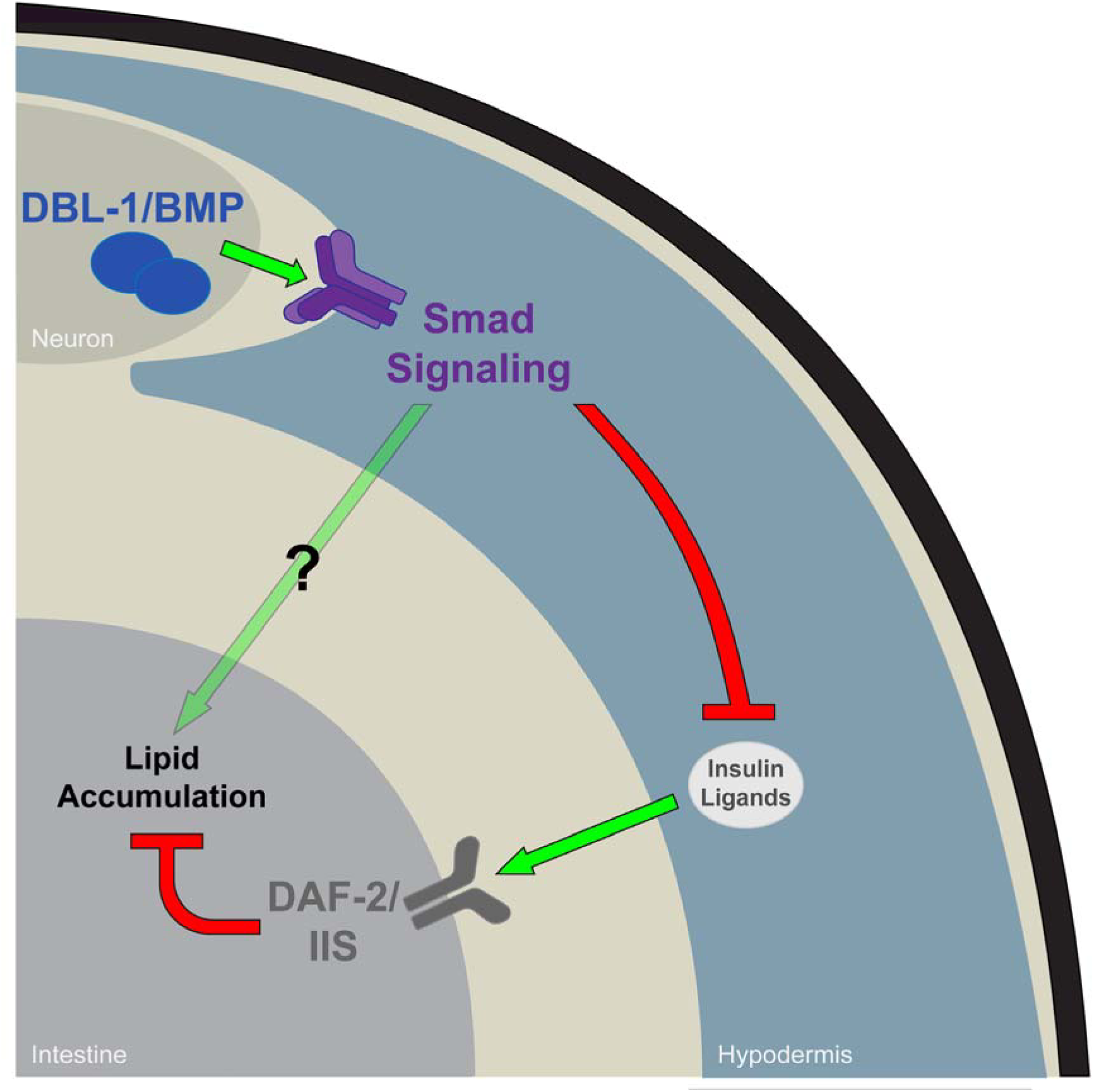
Non-Cell-Autonomous Interaction of DBL-1 and IIS to Regulate Lipids. Our data suggests a necessity for Smad signaling in the hypodermis in order to maintain proper lipid stores. This effect may occur through controlled expression of insulin-like ligands, which in turn regulate the DAF-2/IIS response in the intestine. However, complete epistasis is not observed, indicating an effect on lipid accumulation independent of IIS.

We find that the lipid and body size phenotypes of DBL-1 are separable. Additionally, DAF-2/InsR regulates body size independently of DBL-1. Interestingly, the effect of *daf-2* on body size is opposite of that seen in many other species. In Drosophila, mutations in *InR/InsR* and *chico/IRS* display reduced body size resulting from decreased cell number and size (Bohni et al. 1999). In spite of this, the direction of the lipid-related phenotypes of IIS in Drosophila and in *C. elegans* is the same (Bohni et al. 1999). In mice, as in Drosophila, deficiency in IGF-I or IGF-II causes stunted growth and significant decreases in body weight (Baker et al. 1993). The reason for the shift in growth-regulating function of IIS in *C. elegans* is unknown.

Our findings are part of an emerging body of evidence for interactions between BMP and IIS pathways. In *C. elegans*, DBL-1 and IIS promote germline proliferation (Qi et al. 2017). The Baumeister lab has shown that the IIS transcription factor DAF-16/FOXO and SMA-3/Smad interact in the hypodermis to regulate this germline function. The DBL-1 and IIS pathways have been also shown to regulate reproductive aging, but in this case they act independently (Luo et al. 2009; Luo et al. 2010), similar to their independent functions in body size regulation. In this study, we report the first evidence of interaction between the DBL-1 and IIS pathways that produce an effect in a somatic tissue, the intestine.

In mammalian systems, BMP signaling has been shown to have an effect on insulin sensitivity in mice. Adipocyte-specific knockout of BMP receptor 1A (BMPR1A) exhibited an improved sensitivity to and a slight decrease in insulin (Schulz et al. 2016). Additionally, mice with elevated levels of BMP-4 displayed higher levels of serum insulin and an increase in aberrant phosphorylation of Insulin Receptor Substrate-1 (IRS-1), which prevents proper phosphorylation of IRS-1 and activation of IIS (Chattopadhyay et al. 2017). Our study provides evidence that this interaction is conserved through metazoans. Utilizing *C. elegans* can provide new insights into how the BMP and IIS pathways interact and coordinate.

*C. elegans* has emerged as a powerful genetic model system in which to identify determinants of fat accumulation (Srinivasan 2015). While *C. elegans* do not have dedicated adipose tissue, new insight into the genetic regulation of lipid droplet dynamics can aid in our understanding of the process of lipid homeostasis. Lipid droplets are vital for the utilization and storage of energy at the cellular level and are highly regulated. The morphology of lipid droplets can be an indicator of the function and well-being of a cell or an organism. In mammals, the morphology and utilization of lipid droplets can have an enormous impact on the function of different cell types. Non-adipocyte lipid droplet morphology is maintained in a vastly different pattern than that of adipocytes (Suzuki et al. 2011).

Lipid droplet size is maintained by the balance of triglyceride production and β-oxidation. When peroxisomal β-oxidation is inhibited, such as in *maoc-1, dhs-28*, and *daf-22* mutants in *C. elegans*, average lipid droplet size is increased significantly over wild type (Zhang et al. 2010). Conversely, when fatty acid synthesis is inhibited, lipid droplet size is decreased. Animals containing mutations in *fat-6* and *fat-7*, genes responsible for the first desaturation step in creating polyunsaturated fatty acids, display a significant decrease in average lipid droplet size (Shi et al. 2013). Multiple nutrient sensing pathways exist to provide regulatory signals that dictate lipid droplet mobilization. Lipid droplet morphology can be closely linked to the input of these pathways as energy demands fluctuate with available food. The lipid droplet size phenotype of *fat-6;fat-7* mutants was shown to be independent of the AMPK and TOR pathways. However, it was shown that *fat-6* and *fat-7* work in conjunction with IIS, as both are required for the large lipid droplet size phenotype of *daf-2/InsR* mutants (Shi et al. 2013).

Our data have introduced DBL-1 signaling as a key factor in maintaining proper lipid droplet homeostasis through IIS. Future work will examine whether DBL-1 signaling influences IIS through the transcriptional regulation of insulin-like ligand genes. As SMA-3 is required in the hypodermis, but not the intestine, it may be that DBL-1 signaling is limiting the production of insulin-like ligands in the hypodermis, reducing the level of DAF-2 signaling in the intestine (Figure 6). We will also examine whether Smads interact with transcription factors downstream of DAF-2/InsR, such as DAF-16/FoxO and SKN-1/Nrf, to regulate lipid metabolism. Finally, the question of how an overexpression of DBL-1 leads to an overall reduction in lipids, while increasing the average diameter of lipid droplets, remains. Addressing these questions promises to yield further insight into whole-body lipid homeostasis by these conserved signaling pathways.

## Materials and Methods

### Nematode strains and growth conditions

*C. elegans* were maintained on *E. coli* (DA837) at 15°C, 20°C, and 25°C as specified. The wild-type strain used in this study was N2. The strains used in this study are as follows: LT121 *dbl-1(wk70*), CB678 *lon-2(e678*), CS24 *sma-3(wk30*), CS152 *sma-3(wk30);qcIs6* [*sma-3p::gfp::sma-3+rol-6*], CS619 *sma-3(wk30);qcIs59 [vha-6p::gfp::sma-3+rol-6*], CS628 *sma-3(wk30);qcIs55 [vha-7p::gfp::sma-3+rol-6*], CS630 *sma-3(wk30);qcIs53* [*myo-3p::gfp::sma-3+rol-6*], BW1490 *ctIs40* [*dbl-1(OE)+sur-5::gfp*], CB1370 *daf-2(e1370*), LIU1 *ldrIs1* [dhs-3p::dhs-3::gfp+unc-76(+)], BX106 *fat-6(tm331*), BX153 *fat-7(wa36*), BX156 *fat-6(tm331);fat-7(wa36*). Crosses were used to obtain: *daf-2(e1370);lon-2(e678), daf-2(e1370);sma-3(wk30*), *dbl-1(wk70);ldrIs1*, *ctIs40;ldrIs1*, *daf-2(e1370);ldrIs1*, *daf-2(e1370);dbl-1(wk70);ldrIs1*.

### Oil Red O Staining

Protocol adapted from (O'Rourke et al. 2009). Animals were collected at the L4 stage in PCR tube caps and washed three times in PBS. Worms were then fixed for 1 hour in 60% isopropanol while rocking at room temperature. The isopropanol was removed and worms were stained overnight with 60% Oil Red O solution while rocking at room temperature. Oil Red O was removed and the worms were washed once with PBS w/ .01% Triton and left in PBS. Worms were mounted and imaged using an AxioCam MRc camera with AxioVision software. Images were taken using a 40X objective. Oil Red O stock solution was made with 0.25g Oil Red O in 50mL isopropanol. Intensity of the post-pharyngeal intestine was determined using ImageJ software. Statistical comparisons (one-way ANOVA and Tukey’s multiple comparisons test) were performed using GraphPad Prism 7 software. For each experiment, n>20 per strain repeated in triplicate.

### Body size

Animals were collected at the L4 stage and anesthetized with sodium azide. The worms were then imaged using a QImaging Retiga EXi camera with QCapture software. The midline of each worm was then measured in ImagePro. Statistical comparisons (one-way ANOVA and Tukey’s multiple comparisons test) were performed using GraphPad Prism 7 software. For each experiment, n>30 per strain repeated in triplicate.

### Electron Microscopy

Animals were fixed and embedded by standard methods (Hall 1995). Fixation and microscopy was performed by the David H Hall lab. Animals were well-fed adults grown at 20°C. Animals were fixed 2 days post-egg laying. N2 n=4, *dbl-1* n=2, *sma-3* n=5, *sma-9* n=3.

### Lipid Droplet Morphology

Animals were collected at the L4 stage and anesthetized with sodium azide. The worms were then imaged using a Zeiss ApoTome with AxioVision software. Images were taken using a 100X objective. The tail region of the worm was imaged and the diameter and count of all visible lipid droplets in a 400μm^2^ area were measured using AxioVision software. Statistical comparisons (one-way ANOVA and Tukey’s multiple comparisons test) were performed using GraphPad Prism 7 software. For each experiment, n>15 per strain repeated in triplicate.

### Pharyngeal Pumping Rate

Pumping rate was counted by the number of contractions of the pharyngeal bulb per 20 seconds. Two counts were made per worm and averaged. n>10 per strain repeated in triplicate. Statistical comparisons (one-way ANOVA and Tukey’s multiple comparisons test) were performed using GraphPad Prism 7 software.

## Acknowledgements

We thank Vanessa Almonte, Allie Dananberg, and Mariya Fabisevich for preliminary fat staining and body size experiments. We thank Hasan Zumrut for performing pharyngeal pumping analyses. We are grateful to Ross Cagan and Alicia Meléndez for critical reading of the manuscript. This work was supported in part by National Institutes of Health R15GM112147 and R15GM097692 to CSD and by National Institutes of Health OD010943 to DHH. Some strains were provided by the CGC, which is funded by NIH Office of Research Infrastructure Programs (P40OD010440). This work was carried out in partial fulfillment of the requirements for the Ph.D. degree from the Graduate Center of City University of New York (JFC).

---

**Figure S1.**
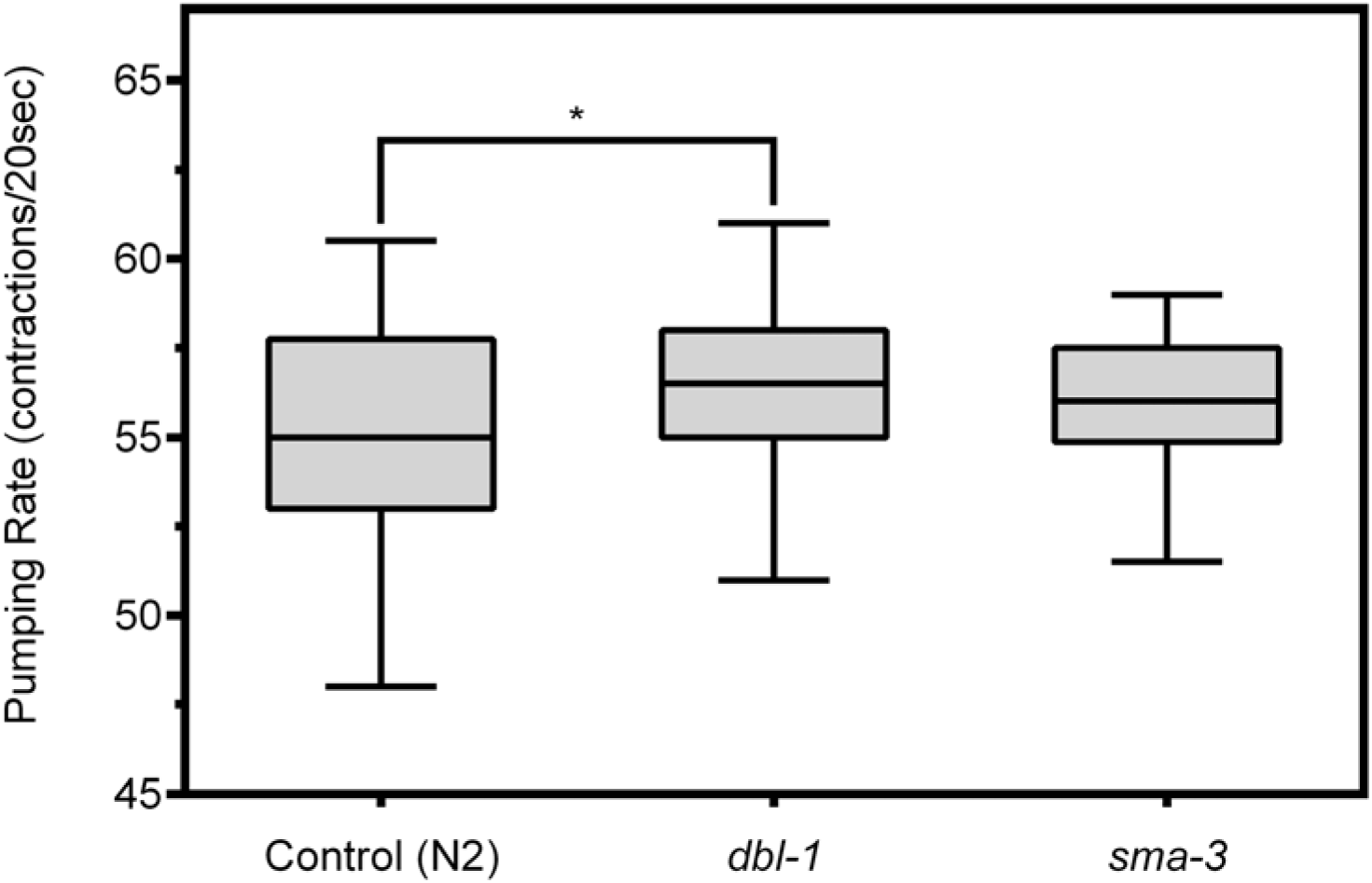
Pharyngeal Pumping Rate in DBL-1 Pathway Mutants. The pharyngeal pumping rate *dbl-1* is slightly higher than that of wild type, increased on average by only 1 extra pump every 20 seconds. *sma-3* animals showed no significant change in pumping rate compared to wild type. DBL-1 pathway mutants appear to eat at the same rate, therefore their lack of lipid stores is not a result of nutrient deprivation. Animals were grown at 20°C. * p value < .05, error bars denote SEM.

